# μ-Lat: A Mouse Model to Evaluate Human Immunodeficiency Virus Eradication Strategies

**DOI:** 10.1101/2020.02.18.955492

**Authors:** Hannah S. Sperber, Padma Priya Togarrati, Kyle A. Raymond, Mohamed S. Bouzidi, Renata Gilfanova, Alan G. Gutierrez, Marcus O. Muench, Satish K. Pillai

**Affiliations:** Vitalant Research Institute, San Francisco, California, United States of America; Free University of Berlin, Institute of Biochemistry, Berlin, Germany; University of California, San Francisco, California, United States of America

**Keywords:** Humanized mouse, HIV, HIV latency, latency reversal, shock and kill, antiviral gene therapy

## Abstract

A critical barrier to the development of a human immunodeficiency virus (HIV) cure is the lack of a scalable animal model that enables robust evaluation of eradication approaches prior to testing in humans. We established a humanized mouse model of latent HIV infection by transplanting “J-Lat” cells, Jurkat cells harboring a latent HIV provirus encoding an enhanced green fluorescent protein (GFP) reporter, into irradiated adult NOD.Cg-*Prkdc*^*scid*^ *Il2rg*^*tm1Wjl*^/SzJ (NSG) mice. J-Lat cells exhibited successful engraftment in several tissues including spleen, bone barrow, peripheral blood, and lung, in line with the diverse natural tissue tropism of HIV. Administration of tumor necrosis factor (TNF)-α, an established HIV latency reversal agent, significantly induced GFP expression in engrafted cells across tissues, reflecting viral reactivation. These data suggest that our murine latency (“μ-Lat”) model enables efficient determination of how effectively viral eradication agents, including latency reversal agents, penetrate and function in diverse anatomical sites harboring HIV *in vivo*.

## Introduction

The advent of antiretroviral therapy (ART) has dramatically reduced morbidity and mortality for human immunodeficiency virus (HIV)-infected individuals with access to healthcare in resource-rich countries. However, despite years of potent therapy, eradication of infection is not achieved due to the persistence of HIV latently-infected cells during treatment(1). Accumulating evidence suggest that “non-AIDS” cardiovascular, renal and hepatic diseases are amplified by HIV infection, and the immune system may exhibit premature senescence even among patients with complete viral suppression(2). Moreover, although enormous progress has been made to provide ART in resource-limited settings, there are huge economic, political and operational challenges to reach the goal of universal access to lifelong treatment. These realities have created a pronounced interest in developing HIV cure strategies.

A number of approaches to achieve a sterilizing or functional cure for HIV infection are currently under investigation, including therapeutic vaccination, immunomodulatory approaches, therapeutic HIV latency reversal (the “shock and kill” strategy), as well as a number of gene therapy approaches(3,4). In all scenarios, it will be critical to have proper diagnostic tools and models in place to comprehensively evaluate performance and safety prior to deployment in a clinical setting. A critical barrier to the development of an HIV cure is the lack of an accessible and scalable preclinical animal model that enables robust evaluation of candidate eradication approaches prior to testing in humans(5). As a result, many promising curative approaches never graduate past the petri dish stage. Infection of nonhuman primates (NHP) with simian immunodeficiency virus (SIV) is an option and has been utilized extensively to study HIV/AIDS pathogenesis(6,7). Recent advancements have been made in optimizing ART regimens to achieve durable virus suppression and thus enable evaluation of HIV cure strategies in the SIV-NHP model(8–10). However, biological limitations remain since this model uses SIV and might not recapitulate human host-HIV interactions and HIV latency mechanisms(11–15). In addition, NHP experiments involve complex ethical considerations, and the high costs and labor requirements only allow small numbers of animals to be utilized in any given trial, limiting statistical power and generalizability(11).

Mouse models represent another alternative with lower costs, more convenient husbandry requirements, as well as greater scalability. In the context of HIV studies, a wide range of small animal models have been developed comprising knockout mouse (16–18), transgenic mouse(19–23), and humanized mouse models(6,24–29). Humanized mice are established by xenotransplantation of human cells or tissues in immunodeficient mouse strains. Most strains used in HIV research are derivatives of severe combined immunodeficient (SCID) mice, which harbor mutations in the gene coding for a DNA-dependent protein kinase catalytic subunit (Prkdc). These mice are “humanized” using two approaches: 1) human cells are injected with or without prior irradiation of mice or 2) portions of tissue are surgically implanted. Different cells have been used for injection, including human peripheral blood mononuclear cells (PBMCs), as in the *hu-PBL-SCID* mouse model(30), obtained from healthy(31) or HIV-infected ART suppressed individuals(32), or human hematopoietic stem cells (HSCs), as in the *hu-HSC* mouse model(33), or the more recently developed *T-cell-only*(34) and *myeloid-only*(35) mouse models (*ToM* and *MoM*). Implantation of fetal thymus and liver tissue fragments are used for the *SCID-hu thy/liv*(36,37) and *bone marrow/liver/thymus* (BLT)(38,39) mouse models. Although all these model systems have contributed to our understanding of HIV pathogenesis and persistence, key limitations remain that need to be addressed in order to fully exploit the potential of these small animal models in HIV cure research. *Hu-PBL-SCID* mice struggle with the development of graft versus host disease (GVHD) which renders this model inapplicable for long-term studies involving HIV persistence. The generation of *SCID-hu thy/liv* and BLT mice is limited due to the need for surgical implantation and a limited supply of tissue. Moreover, these models as well as *hu-HSC, ToM* and *MoM* mice rely on the engraftment of cells or tissues typically derived from human fetal specimens, which in light of recent changes in U.S. federal policies face significant challenges, as ethical, legal, and political considerations surrounding the use of fetal tissue in scientific research have made it increasingly difficult to obtain such material(40). A major limitation shared by all these models remains the low frequency of HIV latently-infected cells, which impacts the applicability of these models as robust *in vivo* test bases for HIV cure strategies.

In the present study, we therefore pursued a new approach and transplanted J-Lat 11.1 cells (J-Lat cells) into irradiated adult NOD.Cg-*Prkdc*^*scid*^ *Il2rg*^*tm1Wjl*^/SzJ (NSG) mice. J-Lat cell clones have been derived from Jurkat cells, an immortalized human T lymphocyte cell line. Each cell harbors a single latent HIV provirus encoding an enhanced green fluorescent protein (GFP) reporter in place of the *nef* gene(41). The proviruses within the J-Lat clones are typically latent due to epigenetic repression(42) and were selected to be responsive to TNF-α stimulation, resulting in viral LTR-driven GFP expression. J-Lat 11.1 cells were chosen for this model as they displayed the greatest viral reactivation upon TNF-α treatment in cell culture settings with respect to other clones(43). Our data show robust engraftment of J-Lat cells in several tissues three weeks after injection as well as significant reactivation of these latently-infected cells *in vivo* upon intravenous administration of the established latency reversal agent (LRA) tumor necrosis factor (TNF)-α. By applying an established and widely-utilized HIV latency reporter cell line(41,43,43), we circumvent the need for fetal or any other donor-derived tissue, and achieve high frequencies of HIV latently-infected cells *in vivo* on a short experimental time scale. In addition, the presence of the GFP reporter cassette in the integrated viral genome provides for quick and convenient assessment of viral reactivation using flow cytometry or microscopy. Moreover, by taking advantage of an HIV reporter cell line, our murine latency (μ-Lat) model represents a highly reproducible platform that allows for a simple transition from *in vitro* screening approaches to an *in vivo* system enabling the evaluation of bioavailability and pharmacokinetics of candidate regimens. Although not intended to serve as a pathophysiology model, we present this approach as a scalable, accessible, and cost-effective preclinical testbed to evaluate the safety, tolerability, and performance of HIV cure strategies in distinct anatomical niches.

## Materials and Methods

### Cell culture and treatment

J-Lat 11.1 cells (kindly provided by Dr. Eric Verdin) contain an integrated latent full-length HIV genome harboring a mutation in the *env* gene and GFP in place of the *nef* gene as a reporter for transcriptional activity of the provirus(45). J-Lat 11.1 cells were grown in media composed of RPMI 1640 (Gibco) supplemented with 10% fetal bovine serum (FBS) (Corning) and 1% Penicillin-Streptomycin (Gibco). Cells were cultured at 37°C in a humidified incubator containing 5% CO_2_. To test the potency of latency reversal agents (LRAs), 1 × 10^6^ cells in 1 ml RPMI were left untreated (negative control), incubated in 0.5% DMSO (vehicle control), 20 nM PMA/1 μM Ionomycin (positive control) or 20 ng/μl TNF-α for 24h.

### Mice

The work was approved by the Institutional Animal Care and Use Committee guidelines at Covance Laboratories, Inc. (San Carlos, CA) under Animal Welfare Assurance A3367-01 and protocol number IAC 2185 / ANS 2469. Animal husbandry was carried out according to the recommendations in the Guide for the Care and Use of Laboratory Animals of the National Institutes of Health. Mice were sacrificed in accordance with the guidelines from the American Veterinary Medical Association.

Adult female and male mice (≥8 weeks old) were included in this study and were maintained at Vitalant Research Institute (VRI). Breeding pairs of NSG mice were obtained from Jackson Laboratories (Bar Harbor, ME), and were bred and maintained at VRI in a vivarium free from >40 murine pathogens as determined through biannual nucleic acid testing (Mouse Surveillance Plus PRIA; Charles River) of sentinel mice exposed to mixed bedding. Mice were maintained in sterile, disposable microisolator cages (Innovive, Inc.), which were changed every 14 days.

Environmental enrichment was provided by autoclaved cotton Nestlets (Ancare Corp.) and GLP-certified Bio-Huts (Bio-Serv). Feed consisted of sterile, irradiated diet of Teklad Global 19% protein diet (Envigo) with free access to sterile-filtered, acidified water (Innovive, Inc.). Several days prior to radiation and 3 weeks following radiation, mice were fed with irradiated Global 2018 rodent diet with 4100 ppm Uniprim® (Envigo).

### J-Lat cell surface marker staining

For each staining, 1 × 10^6^ J-Lat cells were washed once with PBS (Gibco), resuspended in 100 μl PBS, and stained with Zombie dye (Cat. #423105, BioLegend) according to manufacturer’s protocol (1:100 dilution) to enable subsequent discrimination between live and dead cells. 10 min after incubation with the Zombie dye, human CD45-PE (clone HI30, Cat. #304039, BioLegend), CD4-PE (clone OKT4, Cat. #317410, BioLegend), TCR α/β-PE (clone IP26, Cat. #306708, BioLegend), CD27-PE (clone M-T271, Cat. #356406, BioLegend), CD147-APC (clone HIM6, Cat. #306214, BioLegend), CD29-APC (clone TS2/16, Cat. #303008, BioLegend), and HLA-ABC-APC/Cy7 (clone W6/32, Cat. #311426, BioLegend) antibodies were added respectively, and were incubated for another 20-30 min at room temperature (RT) in the dark. Cells were then washed with 2 ml of cell staining buffer (Cat. #420201, BioLegend), resuspended in 300 μl PBS and measured using an LSR II flow cytometer (BD Biosciences). Respective Isotype stained samples were used as control. At least 10,000 events were recorded for each sample.

### J-Lat cell transplantation into mice

If not otherwise stated, J-Lat cell transplantation was performed as follows: mice were irradiated with 175 cGy radiation dose using a RS2000 X-Ray Biological Irradiator (RAD Source Technologies, Inc.) 3 hours prior to cell transplantation. Mice were transplanted with 10 × 10^6^ J-Lat cells in a volume of 200 μl by tail vein injection. J-Lat cell transplanted mice as well as control mice (left untransplanted) were sacrificed and analyzed within 25 days of cell transplantation.

### Engraftment analysis of J-Lat cells in mice by flow cytometry

Transplanted mice were sacrificed by cervical dislocation 3 weeks post injection (or as indicated in figure legends). In one experiment, BM, brain, gut, intraepithelial lymphocytes (IEL) from gut tissue, heart, spleen, lung, lymph nodes, and PB were harvested. In subsequent experiments, selected tissues were harvested including BM, spleen, lung, and PB. Single cell suspensions from the BM, spleen, and PB were prepared as described previously by Beyer et al.(46). Brain, gut, heart, lymph node, and lung specimens were processed as follows: after harvest, all samples were stored in PBS on ice. Brain and lymph node specimens were washed twice in PBS, brain specimens additionally cut into small fragments (3-4 mm^2^), followed by mashing the tissue fragments with a pestle and passaging the cell suspensions through 70 μm cell strainers. Gut, heart, and lung specimens were washed twice with PBS and then cut into 3-4 mm pieces in a petri dish in 1 ml digestion solution consisting of 1 mg/ml DE Collagenase (Cat. #011-1040, VitaCyte) and 100 U/ml DNase I (Cat. #D5025-15KU, Sigma-Aldrich) final concentration in HBSS (Gibco). Fragments were transferred to 50 ml falcon tubes, digestion solution was added to a final volume of up to 5 ml, and samples were incubated at 37°C for 30 min (gut, heart) or 50 min (lung). IEL from gut tissue were isolated as supernatant after digestion incubation. Afterwards, up to 5 ml of stop solution, consisting of 0.5% BSA (Cat. #A2153, MilliPoreSigma) and 2 mM EDTA (Cat. #E0306, Teknova) final concentration in PBS, was added to each sample to end the enzymatic digestion reaction. Single cell suspensions were prepared by passage through a 70 μm cell strainer and washed with PBS. For the analysis of J-Lat cell engraftment, single cells from harvested mouse tissues were stained with human CD147-APC or human CD147-APC combined with human CD29-APC, and additional Pacific Blue conjugate, mouse-specific lineage markers including CD45-Pacific Blue (clone 30-F11, Cat. #103126, BioLegend), TER-119-Pacific Blue (clone TER-119, Cat. #116232, BioLegend), and H-2K^D^ -Pacific Blue antibodies (clone SF1-1.1, Cat. #116616, BioLegend). Cells were incubated for 30 min at RT in the dark. Zombie dye staining detected in the APC-Cy7 channel was used for discrimination of live and dead cells. Following staining, cells were washed and run on an LSR II flow cytometer, where at least 10,000 events were recorded per sample. Flow cytometry data were analyzed using FlowJo software (FlowJo, Inc.).

### Treatment of J-Lat engrafted mice with latency reversal agent TNF-α

Mice engrafted with J-Lat cells were treated with recombinant human TNF-α (Cat. #PHC3011, Gibco), a potent and well-known LRA. Briefly, NSG mice were transplanted with J-Lat cells and 3 weeks post-transplantation, mice received TNF-α (diluted in 200 μl PBS) intravenously at a dose of 20 μg/mouse(47). NSG mice injected with J-Lat cells and treated with PBS were used as vehicle control group.

After 24h of TNF-α treatment, mice were sacrificed to determine viral reactivation in PB, BM, lung and spleen tissues. Latency reversal of the HIV provirus was analyzed by comparing GFP expression of J-Lat cells in tissues of TNF-α treated mice vs the vehicle control group. GFP expression of engrafted J-Lat cells was determined by flow cytometry as described above.

### Data analysis

All data were analyzed using GraphPad Prism version 8.2 and are presented as mean ± the standard error of the mean (SEM). Applied statistical tests and analyses are described in figure legends. A P value of ≤0.05 was considered as statistically significant. Individual P values are indicated in figures.

## Results

### The human cell surface proteins CD147 and CD29 enable discrimination of J-Lat cells from mouse cells

Our model involves transplantation of J-Lat cells (immortalized human T cells harboring a latent HIV provirus with a GFP reporter reflecting viral transcriptional activity) into irradiated adult NSG mice(41). We therefore searched for cell surface markers that identify engrafted J-Lat cells in a background of mouse cells, exhibiting three key features: 1) universal expression across J-Lat cells, 2) high per-cell expression on J-Lat cells, and 3) absence of expression on the surface of mouse cells.

Our candidate panel included seven cell surface proteins commonly known to be expressed in human CD4+ T cells: CD45, CD4, TCR α/β, CD27, CD147, CD29, and HLA-ABC. Four of these cell surface proteins were found to be expressed universally among the J-Lat population: CD45, CD147, CD29 and HLA-ABC (**Figure 1A**). The mean fluorescence intensity (MFI) of the four proteins was measured, reflecting the relative abundance of each protein on the cell surface. CD147 exhibited the highest MFI, followed by CD29 (**Figure 1B**). We next tested if the CD29 and CD147 antibodies (both APC conjugated to achieve maximum signal to noise ratio in the APC-channel) showed detectable binding to mouse cells obtained from different tissues. We harvested bone marrow (BM), lung, brain, gut, intraepithelial lymphocytes (IEL) from gut tissue, lymph nodes, heart and spleen from an untransplanted control mouse. In addition, based on Hoggatt et al.(48), who reported that blood parameters such as cell composition significantly varied depending on sampling manner and site, we performed peripheral blood (PB) harvest in three different ways: by retro-orbital bleed (r.-o.), tail vein bleed, or heart bleed. Human CD29 and CD147 staining was combined with antibodies targeting the mouse-specific lineage markers CD45, TER-119, and H-2K^d^ for a positive identification of mouse cells. While 100% of cultured J-Lat cells expressed CD29 and CD147 and were negative for mouse-specific markers (**Figure 1C**), the frequency of CD147^+^ /CD29^+^ cells was negligible or absent across mouse tissues (**Figure 1D and E**). Most tissues harbored no CD147^+^ /CD29^+^ cells, while 0.06% CD147^+^ /CD29^+^ cells were detected in the lung and 1.37% CD147^+^ /CD29^+^ cells were detected in PB upon heart bleed (**Figure 1E**). For both samples, lung and PB heart bleed, these percentages represent a single event detected to be positive for CD147^+^/CD29^+^ (**Figure 1D and E**). Thus, we concluded that among the tested cell surface proteins, human CD147 combined with CD29 function as a specific marker pair for the identification of J-Lat cells engrafted in mouse tissues.

**Figure 1:**
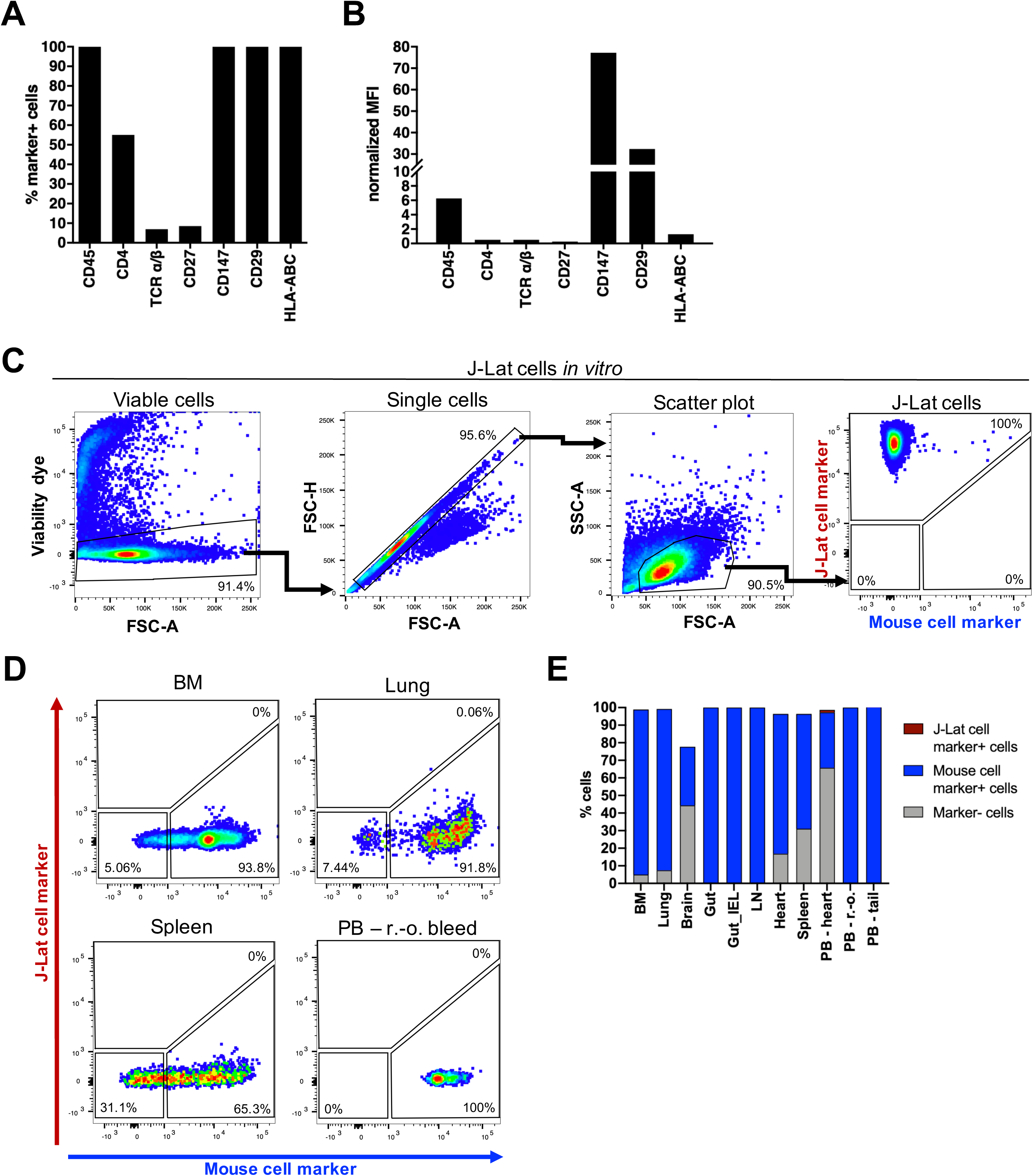
The cell surface proteins CD147 and CD29 are abundantly and specifically expressed in J-Lat 11.1 cells. 1 × 10^6^ J-Lat cells were stained with antibodies targeting select cell surface proteins to (A) measure the frequency of marker-positive cells and (B) mean fluorescence intensity (MFI) of proteins on the cell surface using flow cytometry. The MFI of marker signal positive cells was normalized to signal positive cells of respective isotype control stained samples. A representative experiment is summarized in the bar graphs. (C) Gating strategy to identify human J-Lat cells upon multicolor staining with CD147/CD29-APC (J-Lat cell marker) and CD45/TER-119/H-2Kd-Pacific Blue (mouse cell marker) in vitro. (D) Representative flow cytometry plots show multicolor staining of single cell suspensions prepared from different mouse tissues obtained from an untransplanted control animal. Mouse cells were stained, measured, and analyzed in the same way as *in vitro* J-Lat cells to evaluate specificity and background signal of human CD29 and CD147 antibodies. (E) Single cell suspensions were prepared from 8 mouse tissues and three different harvest methods for peripheral blood (PB) to determine the applicability of the multicolor staining for subsequent engraftment studies. BM = bone marrow, IEL = intraepithelial lymphocytes, LN = lymph node, PB = peripheral blood, r.-o. = retro-orbital.

### Establishment and optimization of J-Lat cell engraftment in humanized mice

In the context of establishing a humanized mouse model, we examined and optimized several parameters simultaneously to achieve a highly reproducible and robust *in vivo* platform (**Figure 2**). In the following pilot experiments, we focused on J-Lat cell engraftment in the BM (**Figure 3**). To measure the kinetics of J-Lat cell engraftment, we harvested the BM at different time points post J-Lat cell injection (**Figure 3A**). We examined the effect of varying cell doses (**Figure 3B**) and irradiation of recipient mice (**Figure 3C**) on J-Lat cell engraftment levels, and compared engraftment levels in NOD.Cg-*Prkdc*^*scid*^ *Il2rg*^*tm1Wj*^*l* Tg(CMV-IL3,CSF2,KITLG)1Eav/MloySzJ (NSG-3GS) and NSG mice (**Figure 3D**). We detected J-Lat cells in the BM 18 days post cell injection (pci), and detection levels reached a significant increase 25 days pci compared to 1 and 3 days pci with an average of 16.3±8.6% engrafted J-Lat cells (**Figure 3A**). Using a J-Lat cell injection dose of 10 × 10^6^ J-Lat cells compared to 5 × 10^6^ J-Lat cells per mouse led to a higher but not significant increase in engraftment levels in the BM of nonirradiated NSG-3GS mice (**Figure 3B**). Nonirradiated NSG-3GS recipient mice harbored lower frequencies of engrafted J-Lat cells in the BM compared to irradiated NSG-3GS recipient mice (**Figure 3C**). Again, this difference was not significant but showed a slight increase in J-Lat cell engraftment levels when recipient mice were irradiated. Finally, when testing engraftment levels in different mouse strains, we found significantly higher numbers of J-Lat cells in NSG mice (**Figure 3D**). J-Lat cell engraftment levels in the BM 3 weeks pci of 10 × 10^6^ J-Lat cells were 2.4 fold higher in NSG mice compared to NSG-3GS mice (**Figure 3D**). Based on our results, we concluded that irradiation of recipient NSG mice and a cell dose of 10 × 10^6^ J-Lat cells per mouse administered through intravenous injection resulted in efficient and reproducible cell engraftment levels which peaked 3 weeks following cell transplantation.

**Figure 2:**
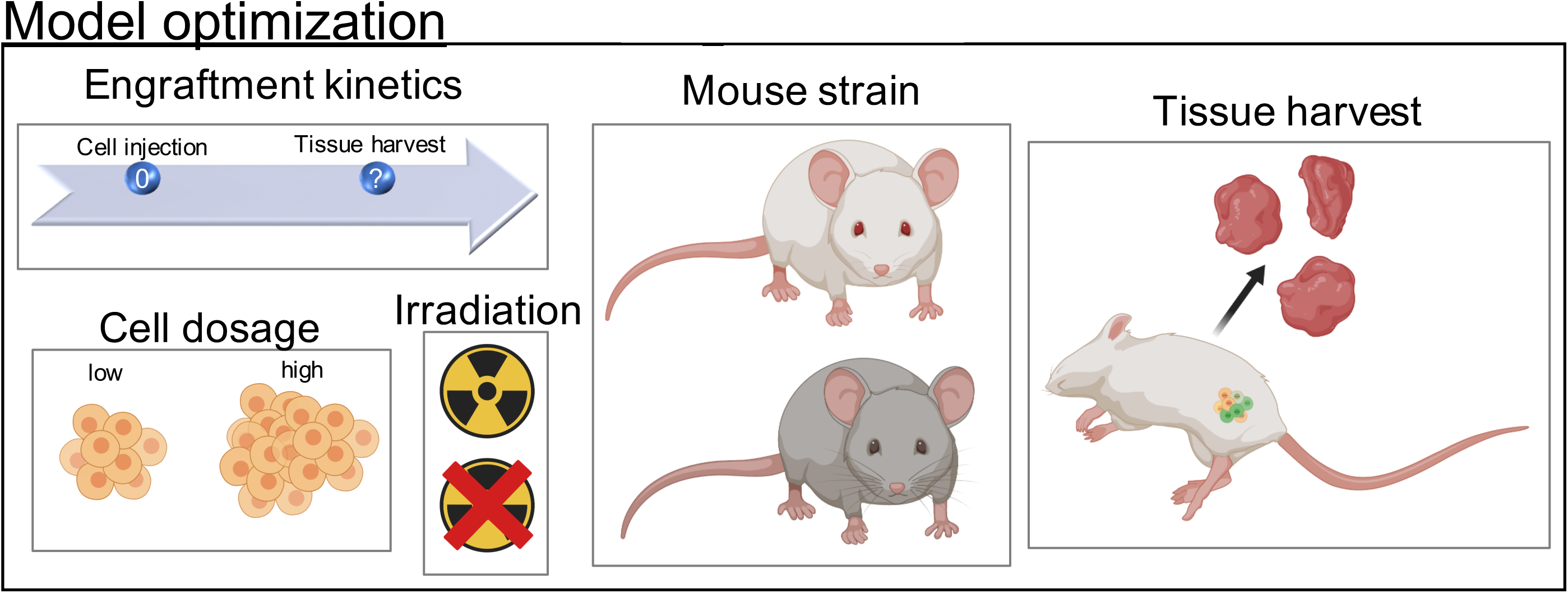
Schematic representation of parameters that were examined and optimized to establish the μ-Lat model. J-Lat cell engraftment kinetics, J-Lat cell dosage, irradiation of recipient mice, and testing of J-Lat cell engraftment in different mouse strains were investigated.

**Figure 3:**
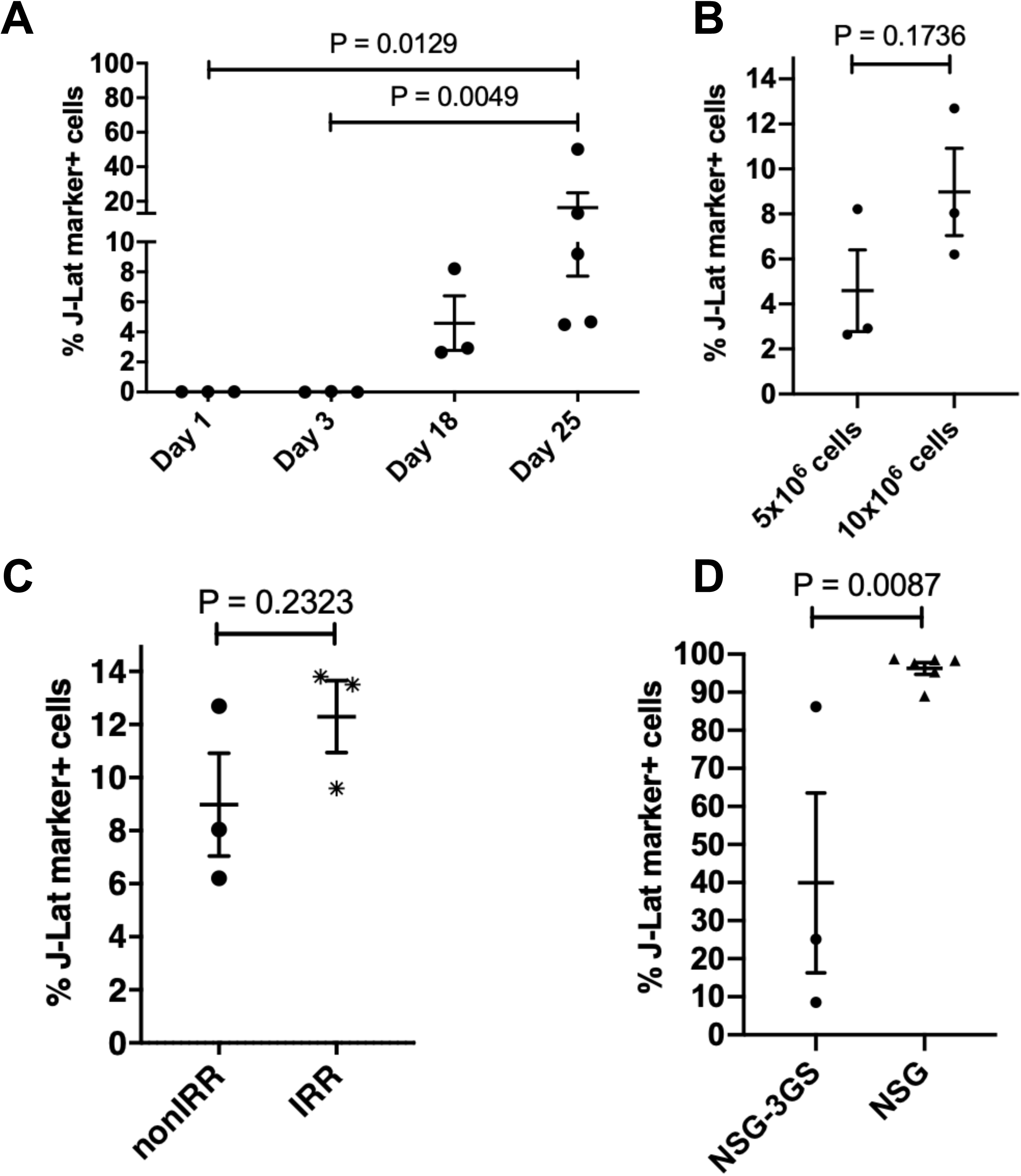
Optimal engraftment levels of J-Lat cells are achieved 3 weeks post injection in irradiated NSG mice using 10 × 10^6^ J-Lat cells for transplantation. The parameters described in **Figure 2** were examined and optimized concomitantly by focusing on engraftment levels in the BM. (A) <u>J-Lat cell engraftment kinetics</u>: Nonirradiated NSG-3GS mice were transplanted with 5 × 10^6^ J-Lat cells and the BM harvested at indicated time points post cell injection. (B) <u>J-Lat cell injection dosage</u>: J-Lat cell engraftment levels in the BM of nonirradiated NSG-3GS were compared using different initial injection cell doses of 5 × 10^6^ J-Lat cells (BM harvested 18 days post cell injection) or 10 × 10^6^ J-Lat cells (BM harvested 14 days post cell injection). (C) <u>Irradiation of recipient mice</u>: NSG-3GS recipient mice were nonirradiated (nonIRR) or irradiated (IRR) before transplantation of 10 × 10^6^ J-Lat cells, and engraftment levels were determined two weeks post cell transplantation. (D) <u>Mouse strain selection</u>: 10 × 10^6^ J-Lat cells were injected either into irradiated NSG-3GS or irradiated NSG mice and engraftment levels in the BM were measured 3 weeks post cell injection and 24h post TNF-α treatment. Each dot represents an individual animal for the respective condition. Each plot represents an independent experiment with n ≥ 3. P values in (A) were determined using the nonparametric Kruskal-Wallis test with the uncorrected Dunn’s test for multiple comparisons. P values in (B-D) were determined using an unpaired, two-tailed t-test. A standard P<0.05 significance threshold was used.

### J-Lat cells engraft successfully in several tissues in transplanted NSG mice

Using the determined experimental conditions, we investigated J-Lat cell engraftment in several tissues, as listed above, that we harvested from 5 irradiated NSG mice 3 weeks pci (**Figure 4**). To analyze the expression of J-Lat and mouse markers on cells obtained from different mouse tissues, we followed the gating strategy as described before (see **Figure 1C and Figure 4A and B**). J-Lat cell engraftment levels were evaluated based on the frequency, MFI, and the absolute number of CD147^+^ /CD29^+^ cells in the respective processed tissues (**Figure 4C-E and Supplementary Figure 1**). We found engrafted J-Lat cells in several tissues including the BM, lung, spleen, and PB (**Figure 4A and B**). The mean frequency of engrafted J-Lat cells varied across tissues from approximately 30% in the BM to 0.7% in the spleen (**Figure 4C**). While the MFI of the engrafted J-Lat cells was comparable within and across tissues (**Figure 4D**), the absolute number of engrafted J-Lat cells varied drastically. We detected the highest J-Lat cell amount in the BM, followed by the amount in the lung and then the spleen (**Figure 4E**). We did not detect engrafted J-Lat cells in the gut, IEL from the gut or heart tissues (**Figure 4C-E**). Processing brain and lymph node tissues yielded only low overall cell numbers that were evaluable (**Supplementary Figure 1A and B**). While J-Lat cells were identified in processed brain tissues at high frequency (**Supplementary Figure 1C**), the actual number of engrafted J-Lat cells was on average 5 cells (**Supplementary Figure 1E**). We thus decided to exclude the brain in subsequent experiments due to the low overall cell numbers. We did not detect engrafted J-Lat cells in the lymph node (**Supplementary Figure 1B-E**). When focusing on engraftment in the PB, we found that heart and r.-o. bleed yielded a substantial higher frequency of engrafted J-Lat cells compared to tail vein bleed (**Figure 4C**). However, PB harvest by r.-o. bleeding showed the highest MFI, and more importantly, the highest absolute number of engrafted J-Lat cells (**Figure 4D and E**). Taken together, we found different engraftment levels based on the sample collection method as observed by Hoggatt et al.(48). Thus, due to considerable engraftment levels and practicality of collection, we decided to harvest PB in subsequent experiments via r.-o. bleeding.

**Figure 4:**
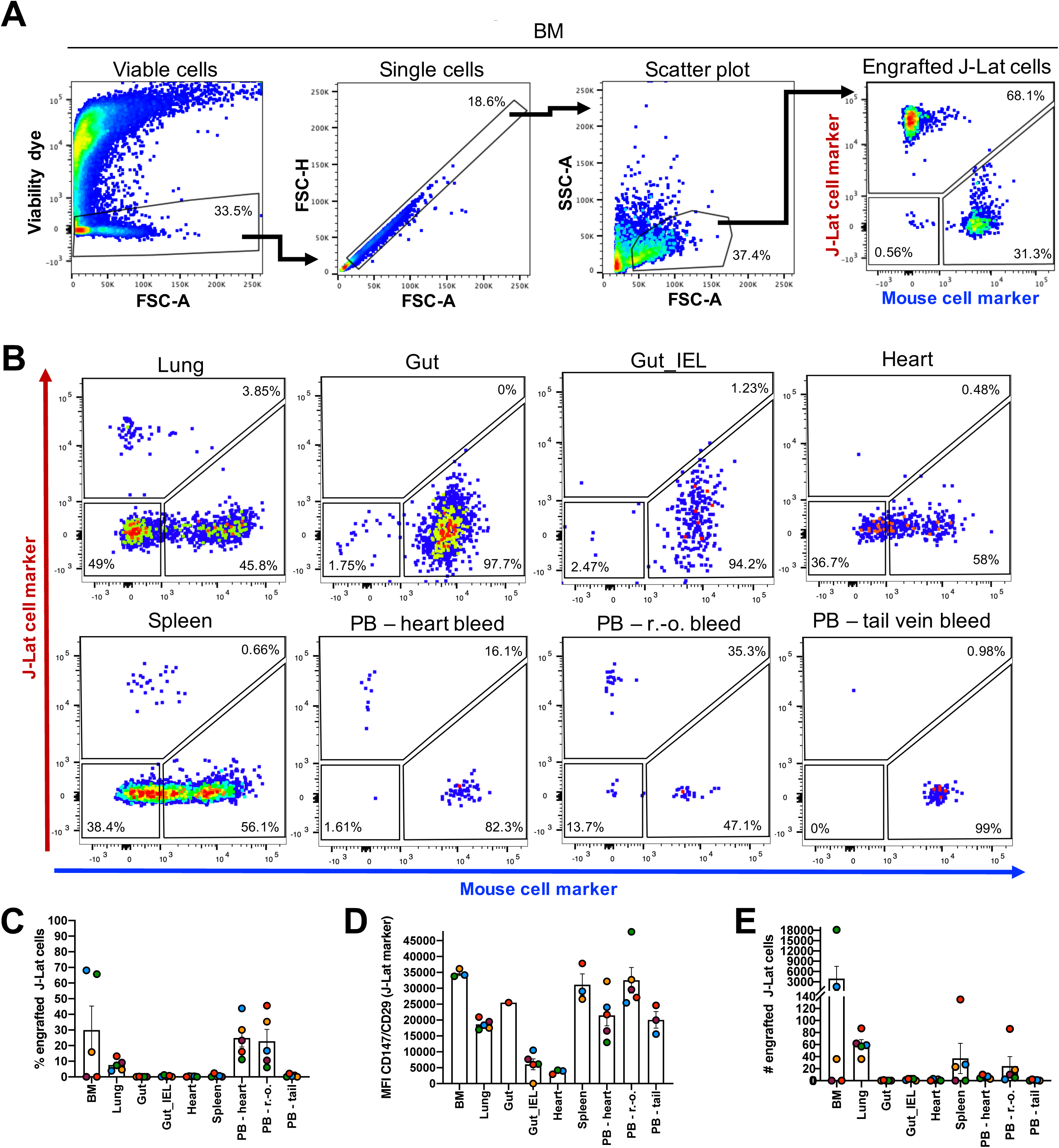
J-Lat cells engraft successfully in several tissues in transplanted NSG mice. (A) Gating strategy to identify human J-Lat cells in harvested mouse tissues exemplified here with BM. (B) Representative flow cytometry plots are shown for engraftment levels across tissue sites observed at necropsy and engraftment levels in PB (n = 5) using three different harvest approaches: r.-o. bleeding, tail vein bleeding, and heart bleeding. Bar graphs summarize (C) frequency, (D) MFI, and (E) number of engrafted J-Lat cells in the respective tissue. Each data point represents an individual animal. Colors indicate tissues harvested from the same animal. Error bars show the standard error of the mean (SEM).

Based on these results and parameter thresholds applied in concert (frequency > 0.005; MFI > 15,000; absolute number > 10), we focused in subsequent experiments on J-Lat engraftment in the BM, lung, spleen and PB via r.-o. bleed and established a suitable engraftment and sample collection protocol to drive the model.

### J-Lat cells maintain low GFP expression upon engraftment in selected tissues

Once we confirmed reproducible and robust engraftment in several tissues, we wanted to investigate GFP expression levels of engrafted J-Lat cells. As described before, the GFP reporter integrated into the latent provirus in J-Lat cells is under the control of the proviral LTR and thus reflects viral transcriptional activity. High GFP expression levels would thus indicate spontaneous viral reactivation *in vivo* and render the model inapplicable for testing LRAs. We measured J-Lat cell engraftment levels and GFP background signal of engrafted J-Lat cells in the selected tissues (BM, lung, spleen, PB r.-o.) of 10 NSG mice 3 weeks pci (**Figure 5**). We applied our previously described gating strategy to identify engrafted J-Lat cells based on CD147/CD29 expression and measured GFP signal within this compartment (**Figure 5A-C**). Engraftment of J-Lat cells was detected in the BM, lung, spleen, and PB r.-o. as observed before (**Figure 5D**). Across these tissues, GFP expression levels of engrafted J-Lat cells remained low and frequencies did not exceed an average of 4% GFP-positive cells (**Figure 5E**). The results showed that while these mouse tissues harbored substantial levels of HIV latently-infected cells *in vivo*, engraftment of J-Lat cells in the tested tissues did not lead to spontaneous reactivation of viral transcription.

**Figure 5:**
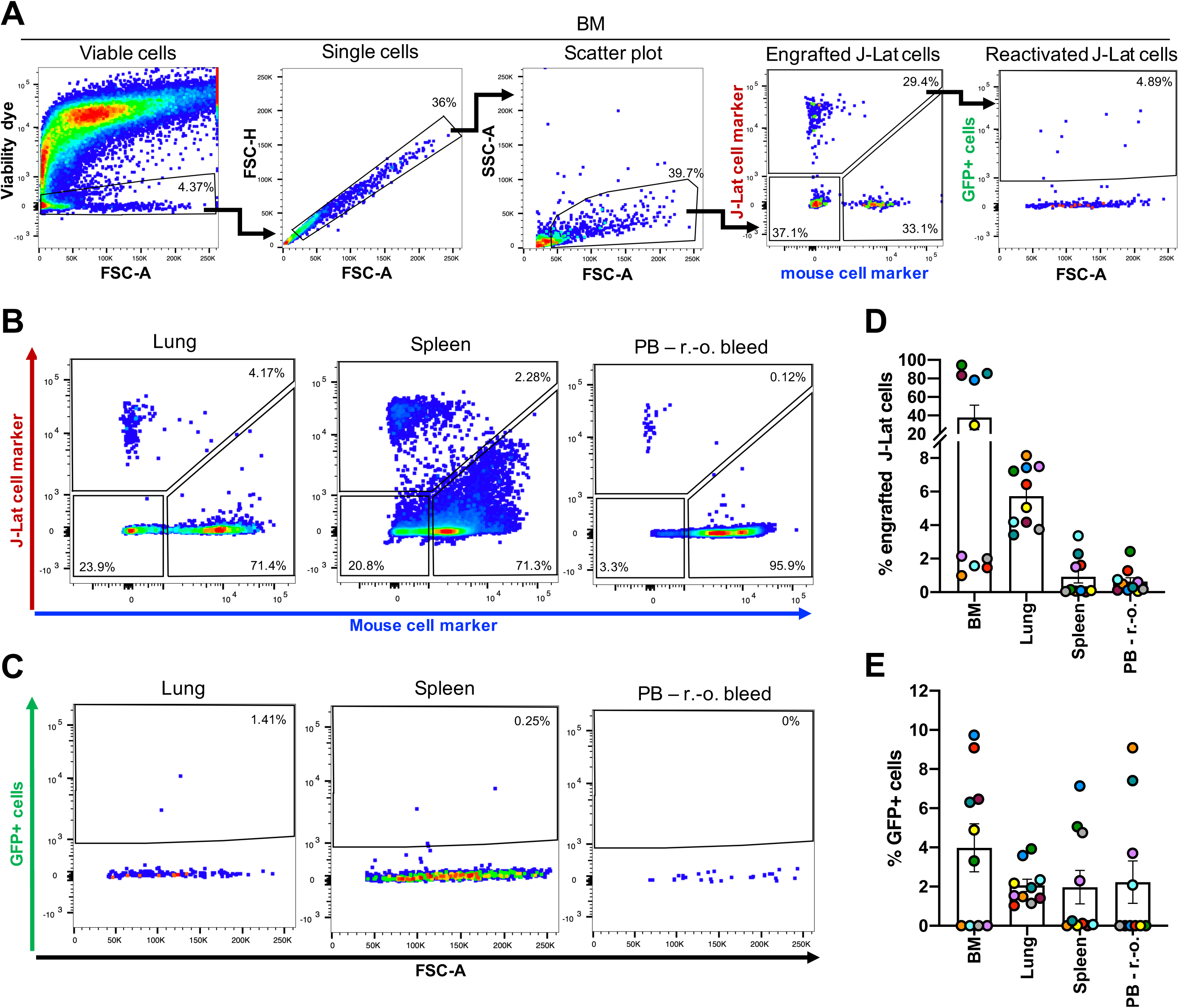
Engrafted J-Lat cells exhibit low basal levels of GFP expression 3 weeks post cell transplantation. (A) Gating strategy to identify human J-Lat cells and GFP background signal of engrafted J-Lat cells in harvested mouse tissues illustrated here with a representative BM sample. Representative flow cytometry plots are shown for (B) engraftment levels and (C) GFP background signal across selected tissues observed at necropsy (n = 10). Bar graphs summarize (D) cell frequency and (E) GFP expression levels of engrafted J-lat cells. Each data point represents an individual animal. Colors indicate tissues harvested from the same animal. Error bars show SEM.

### TNF-α treatment induces GFP expression in tissue engrafted J-Lat cells reflecting reactivation of latent provirus *in vivo*

Finally, we sought to determine if HIV LRA administration would result in viral reactivation *in vivo*. Viral reactivation was measured based on the frequency of GFP expressing cells within engrafted J-Lat cells following LRA treatment (**Figure 6A**).

**Figure 6:**
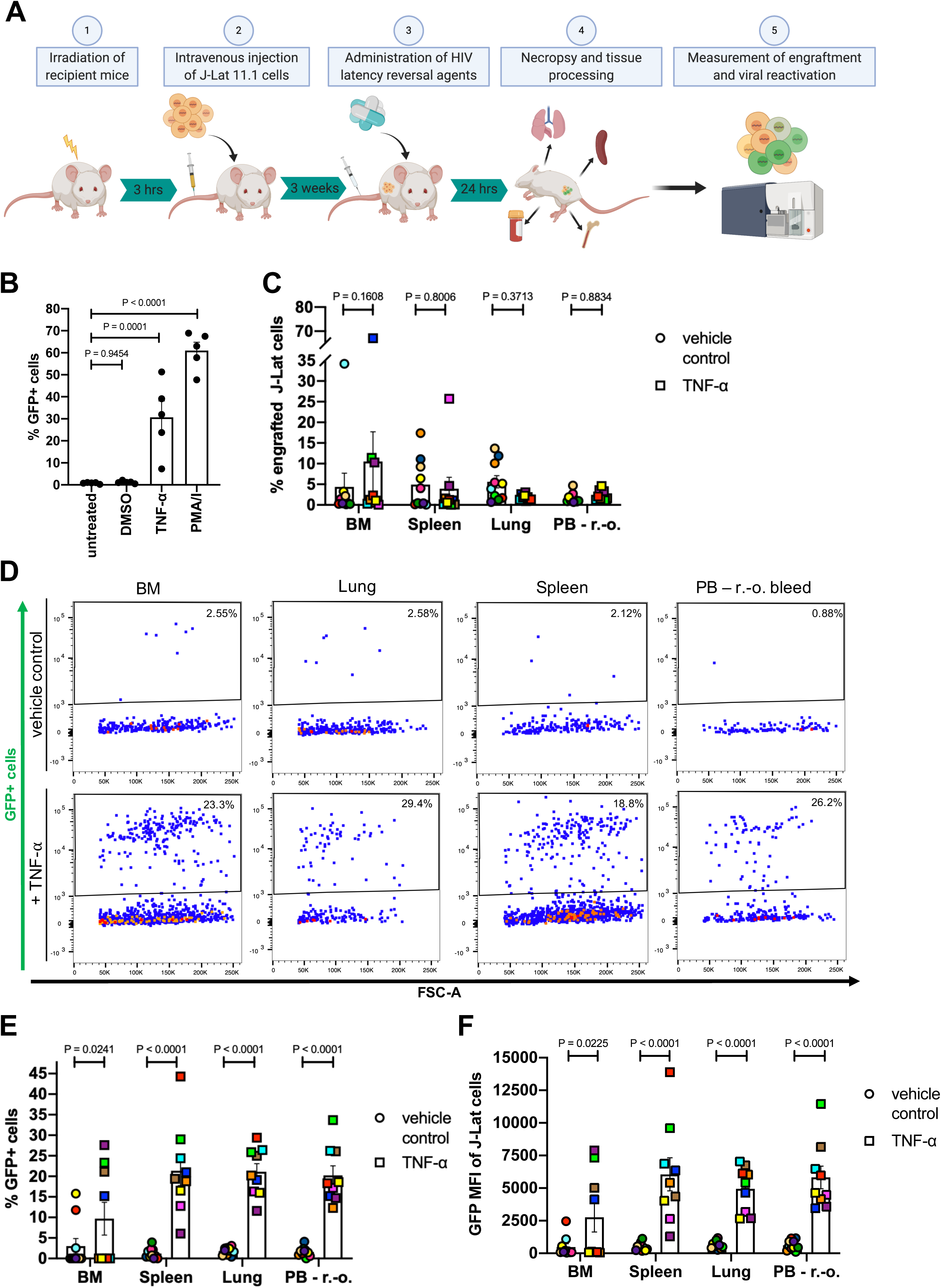
TNF-α treatment reactivates latent HIV *in vivo* in the μ-Lat model. (A) Schematic representation of the procedure to test LRAs *in vivo* using the μ-Lat model: (1) Mice receiving cells of interest for transplantation are irradiated 3 hours prior to cell injection. (2) Each mouse receives 10 × 10^6^ J-Lat cells via intravenous injection. (3) Three weeks post injection (21 days) mice are treated for 24h with LRAs of interest, followed by (4) necropsy, tissue harvest and processing, and preparation of single-cell suspensions. (5) Single-cell suspensions are stained for J-Lat and mouse cell markers to assess engraftment levels via flow cytometry. Reactivation of latent provirus following LRA treatment is assessed by measuring GFP expression via flow cytometry (the J-Lat provirus contains an LTR-driven GFP reporter). (B) J-Lat cells were treated *in vitro* for 24h with 0.5% DMSO (vehicle control), 20 ng/μl TNF-α, 20 nM PMA/1 μM Ionomycin (positive control) or left untreated (negative control) to assess viral reactivation based on GFP expression (n = 5). (C) J-Lat engraftment levels were measured 24h post tail vein injection of 20 μg TNF-α (n = 9) or vehicle control (PBS, n = 10) in BM, spleen, lung and PB of mice. (D) Representative flow cytometry plots demonstrating the effect of vehicle control (top panel) or TNF-α treatment (bottom panel) on viral reactivation (GFP expression) in engrafted J-Lat cells. The induction of GFP expression among engrafted J-Lat cells across tissues is summarized as (E) frequency of GFP+ cells and (F) GFP MFI. One-way ANOVA and uncorrected Fisher’s LSD for multiple comparisons was used to analyze (B) *in vitro* reactivation data, and a mixed effects model and uncorrected Fisher’s LSD for multiple comparisons was used to analyze (C) *in vivo* engraftment and (E-F) viral reactivation data. Colors indicate specific animals and tissues obtained from the same animal. Error bars show SEM.

The proviruses within the J-Lat family of clones were selected to be responsive to TNF-α stimulation, resulting in viral LTR-driven GFP expression(41). Previous studies have shown, that among the established clones, J-Lat 11.1 cells displayed greatest viral reactivation upon TNF-α treatment in cell culture settings(43). 24h *in vitro* treatment of J-Lat cells with 20 ng/μl TNF-α resulted in a significant increase of GFP-positive cells compared to mock-treated (0.5% DMSO) and untreated cells, which were largely GFP-negative (**Figure 6B**). Therefore, we performed *in vivo* reactivation experiments using 20 μg of TNF-α as an LRA (with PBS as a vehicle control)(47). 24h following TNF-α tail vein injection, effects on tissue engraftment and reactivation of latent provirus were estimated in a cross-sectional manner at necropsy, comparing 9 animals treated with TNF-α versus 10 vehicle control animals. J-Lat cell engraftment and GFP expression of J-Lat cells were analyzed as described in **Figure 5A**. Analyses of all tissues demonstrated no significant effect of TNF-α treatment on J-Lat cell engraftment levels within the respective compartment (**Figure 6C**). However, TNF-α treatment did lead to significant increases in the frequency of GFP-positive cells as well as GFP MFI among engrafted J-Lat cells (**Figure 6D-F**). Comparing the two populations (TNF-α vs vehicle control) demonstrated a GFP expression fold-change of 3.2 in the BM, 17.8 in the spleen, 11.7 in the lung, and 12.7 in the PB, illustrating potent and significant viral reactivation.

## Discussion

The development of an HIV cure will be accelerated by the deployment of a convenient and cost-effective animal model that enables the determination of an agent’s therapeutic efficacy as it permeates through tissue-specific barriers and the circulatory, respiratory and excretory systems. We present a novel humanized mouse model of HIV latency that aims to address this gap. In this study, we established a reliable method to discriminate between J-Lat and mouse cells (**Figure 1 and Figure 4**) and demonstrated robust engraftment of these cells in multiple tissues sites (**Figure 4 and Figure 5**). Significant viral reactivation was observed in TNF-α-treated animals with respect to vehicle control-treated animals (**Figure 6D-F**). Importantly, although *in vivo* treatment with TNF-α induced GFP expression and thus viral reactivation by 3.2 (BM), 17.8 (spleen), 11.7 (lung), and 12.7 (PB) -fold compared to vehicle control treated animals (**Figure 6E**), TNF-α treatment *in vitro* induced a 25.3-fold increase in GFP expressing cells (**Figure 6B**). Moreover, the observed variability in TNF-α-mediated viral reactivation across tissue sites demonstrates that the efficacy of latency reversal approaches can differ dramatically between distinct anatomical niches due to pharmacokinetic factors and drug penetration efficiencies. These data emphasize the importance of testing the efficacies of promising and potent regimens in an *in vivo* system, and reinforce the relevance of the μ-Lat model.

Although not intended to serve as a pathophysiology model, the μ-Lat platform offers key advantages over existing preclinical models focused on HIV eradication. Firstly, the model is highly efficient; the timeframe to generate mouse colonies that are ready for administration of experimental therapies is approximately three weeks. In comparison, even the simplified humanized mouse model introduced by Kim et al.(31) requires four weeks for robust engraftment of intraperitoneally injected human PBMCs in NSG mice and an additional 5 weeks of incubation upon HIV infection. Secondly, the presence of the GFP reporter cassette in the integrated J-Lat viral genome allows for rapid and convenient assessment of HIV latency reversal *in vivo*, eliminating the requirement for PCR or culture-based diagnostics to determine LRA potency. Thirdly, the model is characterized by high frequencies of HIV latently-infected cells (e.g. in some experiments exceeding 30% engraftment in BM), and these cells are distributed in a number of relevant anatomical sites that are central to HIV persistence. This stands in contrast to organoid-based models (e.g. the SCID-hu *thy/liv* mouse model) that exclusively involves viral colonization within a xenografted tissue(49). Lastly, the model is highly scalable due to the low costs and limited labor requirements of the method involving the easily and widely available J-Lat cell line (provided free of charge via the NIH AIDS Reagent Program). Importantly, the clonal nature of cell lines promotes consistency and high reproducibility across laboratories, while allowing for an easy transition from the petri-dish stage to an *in vivo* evaluation system.

Similar to other animal models, there are caveats associated with the μ-Lat model. Although the GFP reporter present within the J-Lat cell conveniently broadcasts transactivation of the viral LTR and induction of viral transcription (informative precursor steps that may lead to the production of viral antigen and release of virus), other assays will need to be applied to the μ-Lat model to specifically quantify viral production or clearance of infected cells mediated by a candidate HIV eradication agent. In addition, the efficiency of the model is largely driven by the administration of an HIV latently-infected cell line, rather than differentiated primary cells. As presented here, a single latent clone with a single proviral integration site was injected into mice. Proviral integration site is known to affect viral latency and responsiveness to LRA administration(4,50–50). However, this issue could be ameliorated by mixing different HIV latently-infected cell clones at specific ratios, thereby achieving higher integration site heterogeneity to more accurately represent *in vivo* variability. For instance, there are 11 J-Lat clones currently available via the NIH AIDS Reagent Program, each of which is characterized by a distinct proviral integration site. These clones often behave differently from each other as well when exposed to LRAs in vitro, likely reflecting a diversity of molecular mechanisms reinforcing viral latency across clones(43). This mechanistic diversity could further enhance the predictive potential of the μ-Lat model.

Beyond concerns regarding integration site heterogeneity, the immortalized nature of the J-Lat cell line may impact molecular and regulatory pathways that affect HIV latency. However, the widespread usage of the J-Lat model and its derivative clones to examine viral latency and to evaluate the efficacy of HIV cure strategies *in vitro* speak to the model’s utility(53–62). Ample data from primary cell-based models of latency and experiments involving ex vivo administration of LRAs to cells obtained from HIV-infected individuals on ART suggest that differences between applied models and between individuals can have dramatic effects on the establishment, maintenance, and reversal of HIV latency(43,63,63). Moreover, profiling of HIV latency in multiple tissue sites has demonstrated that the nature of viral latency may even vary extensively within a single infected individual(65–67). Therefore, it needs to be considered that any applied model system or cells from any given individual may introduce biases that impact generalizability, just as a cell line-based system(26).

Our work described here represents a proof of concept, demonstrating that the engraftment of a cell line-based model of HIV latency may constitute a useful testbed for HIV cure strategies. This general approach is highly versatile and should allow for a broad range of infected cell types to be examined *in vivo*. For example, HIV latency in the myeloid compartment is likely critical to viral persistence(35,68–68), and multiple reports suggest this compartment may respond quite differently to curative approaches, as compared to lymphoid reservoirs(72,73). It may be fruitful to inject the U1 HIV chronically-infected promonocytic cell line(74) into NSG mice to examine LRA responses in myeloid cells. Building further on the myeloid theme, the central nervous system (CNS) compartment is an important viral sanctuary site in the setting of ART(73,75–75), and HIV eradication approaches will almost certainly face unique challenges in this niche(78,79). As direct injection of human cells into the murine CNS has been used successfully as an engraftment approach(80–84), site-specific injection of the recently developed HC69.5 HIV latently-infected microglial cell line(85,86) into the brains of NSG mice may provide a convenient platform to gauge CNS-focused cure strategies.

Beyond preclinical investigation of LRAs, the μ-Lat framework may provide a convenient model system to evaluate gene therapy-based HIV eradication approaches. Gene therapy approaches targeting HIV infection generally fall into two categories: 1) Gene editing can be used to target or “excise” the HIV provirus directly in infected cells as an eradication approach(87,88) or 2) Editing can be used to modulate host cells to render them refractory to HIV infection and/or potentiate antiviral immune responses(89). In the former case, *in vivo* delivery of the gene therapy modality will likely be necessary to pervasively attack the HIV reservoir within a broad range of tissue sites in infected individuals. The μ-Lat model is well-suited to investigate gene delivery in this context, as it is characterized by robust engraftment of HIV latently-infected cells into diverse anatomical niches. This will allow for efficient assessment of gene therapy vector dissemination and antiviral function across anatomic sites. The LTR-driven GFP cassette within the J-Lat integrated provirus may facilitate this assessment; direct targeting of the GFP sequence may be used to examine vector trafficking and delivery, while specific targeting of the HIV LTR as a cure approach would be associated with a convenient readout (relative loss of GFP expression upon induced latency reversal).

In summary, the μ-Lat model is optimized for efficient and scalable evaluation of select HIV eradication approaches *in vivo*, allowing determination of therapeutic efficacy in addition to essential safety, tolerability, and pharmacokinetic parameters. Further development and diversification of the μ-Lat model system may enable convenient testing of HIV eradication approaches, including antiviral gene therapy strategies, in a range of cell types and tissue sites *in vivo*.

## Nonstandard abbreviations

ART: antiretroviral therapy
BM: bone marrow
GFP: green fluorescence reporter
GVHD: graft versus host disease
HIV: human Immunodeficiency virus
HSC: hematopoietic stem cell
IEL: intraepithelial lymphocytes
LN: lymph node
LRA: latency reversal agent
LTR: long terminal repeats
MFI: mean fluorescence intensity
NHP: nonhuman primates
NSG: NOD.Cg-*Prkdc*^*scid*^ *Il2rg*^*tm1Wjl*^/SzJ
NSG-3GS: NOD.Cg-*Prkdc*^*scid*^ *Il2rg*^*tm1Wj*^/Tg(CMV-IL3,CSF2,KITLG)1Eav/MloySzJ
PBMC: peripheral blood mononuclear cell
PMA: phorbol-myristate-acetate
r.-o.: retro-orbital
SCID: Severe combined immunodeficiency
SIV: simian immunodeficiency virus
TNF: tumor necrosis factor

## Acknowledgments

This study was supported by the National Institutes of Health grants R01 AI150449 (SKP) and R01 MH112457 (SKP), and was additionally supported by a grant from the National Institutes of Health, University of California San Francisco-Gladstone Institute of Virology & Immunology Center for AIDS Research (P30 AI027763). We thank Dr. Roland Schwarzer for valuable input and careful review of the manuscript.

## Author contributions

HSS, PPT, KAR, MSB conceived and performed experiments; PPT, RG, AGG, MOM performed *in vivo* experiments; HSS, PPT collected and analyzed the data; HSS, PPT, MOM, SKP designed the study and wrote the manuscript.

