## Supplementary Figure 1 for "μ-Lat: A Mouse Model to Evaluate Human Immunodeficiency Virus Eradication Strategies"

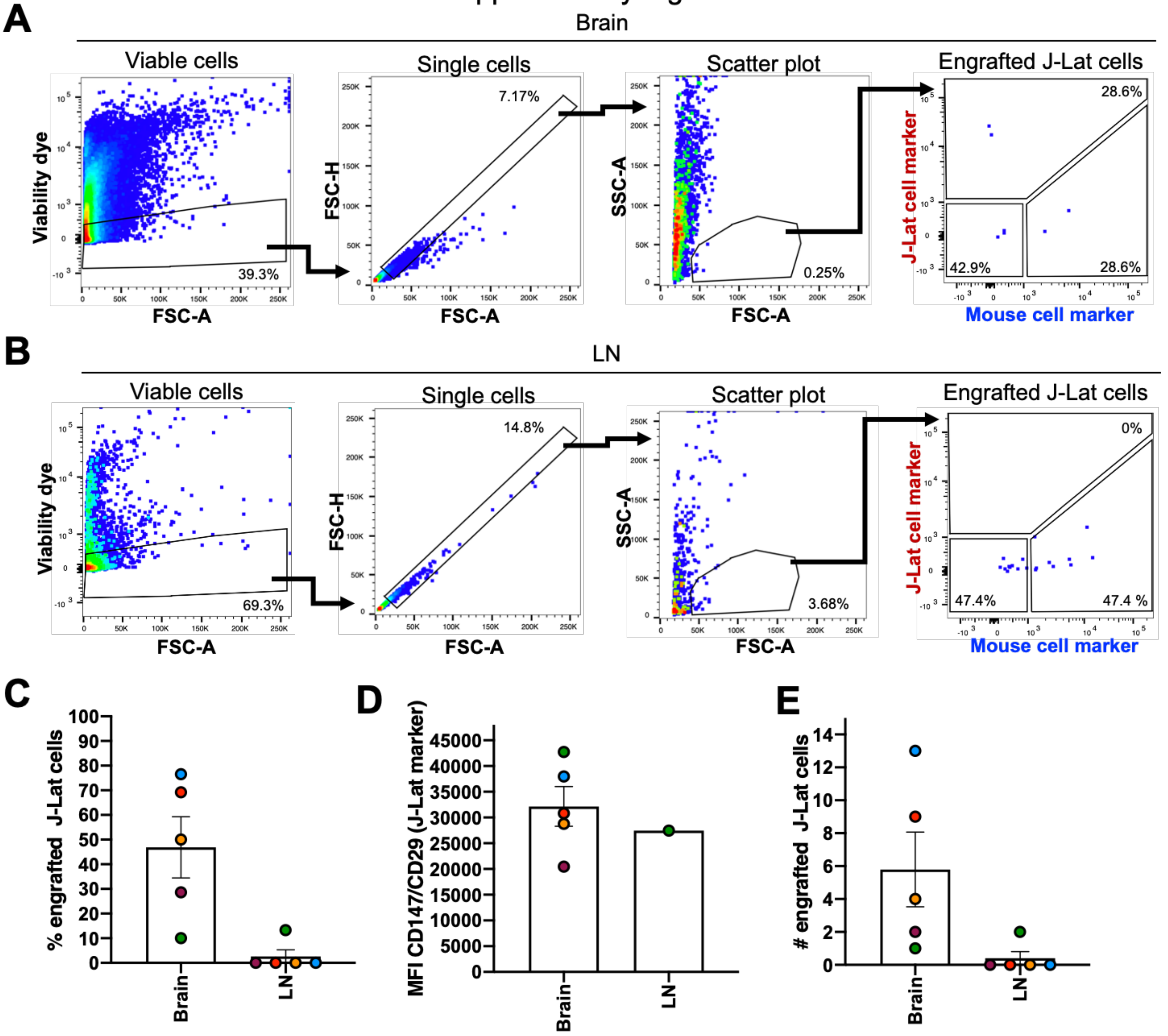


**Supplementary Figure 1: Engraftment of J-Lat cells in brain and lymph node tissues.** Representative flow cytometry plots demonstrating the gating strategy to identify human J-Lat cells in harvested mouse (A) brain and (B) LN tissues (the same gating strategy was applied to all tissues). Bar graphs summarize (C) engrafted J-Lat cell frequency, (D) MFI of engrafted J-Lat cells, and (E) number of engrafted J-Lat cells in the brain and lymph node tissues harvested from 5 NSG mice 3 weeks post cell injection. Each data point represents an individual animal. Colors indicate tissues harvested from the same animal. Error bars show the standard error of the mean (SEM). As displayed in the flow cytometry plots on the right, very low overall cell numbers from brain and lymph node could be analyzed, limiting the interpretability of data describing engraftment levels of J-Lat cells in these tissues.
